# Comparison of Ferroptosis Related Genes expression in Human Oral Squamous Cell Carcinoma and Normal oral tissues

**DOI:** 10.1101/2021.08.08.455486

**Authors:** Rooban Thavarajah, Elizabeth Joshua, Kannan Ranganathan

## Abstract

**Introduction:** Evasion of programmed cell death (PCD) is a hall mark of oncogenesis. There are different types of PCD. Iron related PCD, ferroptosis is being increasingly associated with neoplastic process. There are very few reports that investigate the role of ferroptosis in Oral Squamous Cell Carcinoma (OSCC). An attempt is made to compare the ferroptosis related genes(FRGs) expression in human OSCC and normal oral tissues.

**Materials and Methods:** Gene Expression Omnibus repository was scanned for OSCC mRNA datasets along with normal control tissues. Datasets fulfilling inclusion and exclusion criteria as well as that fulfilled the statistical correlation requirements were considered for this study. Differentially expressed mRNAs were identified. From the literature and ferroptosis database, FRGs were identified and those FRGs were differentially expressed were validated using The Human Cancer Genome dataset.

**Results:** In all 44 FRGs were identified to be differentially expressed between OSCC and control tissues. Of the 44, 21 were that promoted ferroptosis including 18 drivers of ferroptosis. Of the 21 FRGs that drives ferroptosis, 9 were found significantly elevated in controls while the remaining 12 were elevated in OSCC. The role of the differentially expressed FRGs were also studied. Of the 44 FRGs, 36 were validated using the human cancer genome dataset.

**Discussion and Conclusion:** Drivers and suppressors of ferroptosis were differentially expressed in OSCC and controls. This reflects that ferroptosis has a dual role in oncogenesis – both as a promoter and a suppressor. The identified specific FRGs in this studied would help to understand the role of PCD in OSCC progression and help in designing better treatment.

## 1. BACKGROUND

Oral Squamous Cell Carcinoma (OSCC) is one of the common tobacco associated cancer that poses significant treatment challenge. The molecular pathogenesis of OSCC involves several gene level changes, and biological pathways.^[1]^ Foremost of them is the evasion of programed cell death (PCD). PCD is orchestrated by intracellular signal transduction pathways. Based on the morphological features and molecular mechanisms, PCD is divided into apoptotic cell death (where it retains cell membrane integrity and happens in a caspase□dependent manner) and the non□apoptotic cell death (through membrane rupture and caspase□independent manner).^[2]^ The nomenclature committee on cell death listed several non-apoptotic PCD based on involved mechanisms at gene level.^[3]^ Of the several PCDs, ferroptosis is an iron-dependent PCD that employs phospholipid peroxidation and reactive oxygen species (ROS) to cause cell death. Essentially, this is driven by a redox imbalance between the production of oxidants and antioxidants. This imbalance arises from the sum of total abnormal expression and activity of multiple redox-active enzymes.^[3–5]^

Ferroptosis is involved in death-balance in normal cells and tissues.^[6]^ There is an emerging interest in the role of ferroptosis in several pathologies including cancer, with several studies elucidating the multiple pathways and intricate mechanisms of ferroptosis. Certain carcinogenic pathways have been known to modulate key ferroptosis factors causing evasion or induction of ferroptosis in neoplastic cells.^[7–12]^

Ferroptosis can occur via canonical (inactivation of the central antioxidant system) and non-canonical (Fe^2+^ catalysed ROS overload) pathways. The canonical pathway is caused by directly inhibiting glutathione-dependent peroxidase (GPX4) or by causing glutathione (GSH) depletion. The non-canonical pathway involves by increase in the intracellular iron pool. This intracellular free Fe^2+^ mediates formation of lipid hydroperoxide via non-enzymatic lipid autoxidation, enzymatic lipid peroxidation, Fenton reaction, ferratinophagy. This free iron also serve as cofactor for lipoxygenase (LOX) to enzymatically catalyse polyunsaturated fatty acid peroxidation. Metabolically, there are non-enzymatic pathways or the extrinsic or transporter-dependent pathway (e.g., decreased cysteine or glutamine uptake and increased iron uptake) and the intrinsic, enzymatic pathway (eg. GPX4 inhibition). The later includes the lipid metabolism pathway(via ACSL4, non-canonical), FSP1-CoQ10-NADPH pathway (non-canonical), Mevalonate pathway(canonical), GPX4 pathway(canonical), glutaminolysis pathway (canonical), cystine deprivation-induced (via system X_c_^−^, canonical) ferroptosis pathways, iron metabolism pathway (non-canonical) and ferritinophagy(non-canonical).^[7–12]^ Detailed mechanisms of ferroptosis and associated pathway have been elucidated and several genes associated with ferroptosis have been described. As of the date, around 259 genes have been associated with ferroptosis.^[13]^

In a bid to keep a check on neoplastic progress, cells have evolved tumor suppressing pathways and ferroptosis is one among them. For ferroptosis to occur in cancer cells, two central biochemical events - intracellular iron accumulation and lipid peroxidation are required. This happen either by metabolite or non-metabolite mediated ways. The former include diminishing cysteine uptake through the inhibition of system X_C_^−^ and targeting GPX4, resulting in increasing iron concentration and ROS. Depending on cellular state, p53 can inhibit or promote ferroptosis. Ferroptosis is induced by preventing the transcription of the SLC7A11 gene. SLC7A11 encodes for the substrate-specific subunit of system X_c_^−^. Suppression of SLC7A11 effectively blocks cysteine uptake into the cell and suppresses GPX4 activity. Subsequently neoplastic cells undergo ferroptosis upon routine oxidant insults. Cells to induce ferroptosis may also activate lipoxygenase ALOX12. The transcriptional repression of SLC7A11 leads to ALOX12-dependent ferroptosis. In a similar fashion, other lipoxygenases such as ALOXE3 and ALOX15B mediate ferroptosis in cancer.^[7–12]^

The p53-p21 axis, preserves GSH, thiols, and suppresses phospholipid oxidation, p53 which can reduce ROS, thereby having an anti-ferroptotic effect. It also modulates the mevalonate pathway via ATP-binding cassette subfamily A member 1 (ABCA1) and sterol regulatory element binding protein 2 (SREBP2) to function as an anti-ferroptotic mediator.^[6,14]^

In normal and pathological conditions, ferroptosis is precisely regulated and orchestrated at multiple levels, including epigenetic, transcriptional, posttranscriptional and posttranslational pathways. Epigenetic regulation also plays a definitive role in ferroptosis. For example, the loss of function mutations of the tumor suppressor BRCA1-associated protein-1(BAP1) is associated with ferroptosis. In addition, many long non-coding RNAs and microRNAs have been proven to be associated with ferroptosis through ROS.^[6]^

The association of ferroptosis related genes (FRG) with OSCC have not been widely reported.^[15–17]^ The aim of this manuscript is to identify the association of FRG by comparing the gene expression of OSCC with normal oral tissues.

## 2. MATERIALS AND METHODS

### 2.1 Source of microarray datasets

A systematic search of the National Centre for Biotechnology Information’s Gene Expression Omnibus (GEO) repository (NCBI-GEO - http://www.ncbi.nlm.nih.gov/geo) using the keywords “Oral cancer” “OSCC” Oral Squamous cell carcinoma” was carried out in early May 2021, without any restriction of age of the dataset. The microarray datasets and their descriptions contents were further search-limited to

- Humans - *Homo sapiens*
- Type - “expression profiling by the array”
- Recommended or acceptable, extraction protocol
- Has mRNA.
- Ideally had matched control oral cavity tissue, but if study lacks control tissues, it was still considered for “OSCC” group

Datasets that had unclear extraction protocols or sources or details were excluded. No emphasis was placed on types of dataset platforms. The gene series were collated and characterized to have pooled “OSCC” and matched OSCC free “controls”

### 2.2 Pair Wise Comparison

The individual patient’s gene expression datasets captured by methodically reviewing and screening the microarray datasets. They were collated using the pair wise comparison of the ExAtlas online web tool at https://lgsun.irp.nia.nih.gov/exatlas.^[18]^ The quantified genes were then log_2_ transformed, normalized with quantile normalization method and later all datasets combined. Principal component analysis (PCA) was performed to check the tissue/gene distribution with FDR ≤0.05, correlation threshold = 0.7 and fold change threshold of 2. In the tool, PCA is calculated using the Singular Value Decomposition (SVD) method that generates eigen vectors for rows as well as columns of the log-transformed data matrix.^[18]^ For plotting of mRNA of genes (biplot) column projections were used, as biplots is helpful for visually exploring associations between cases and controls.

A pairwise compared with ANOVA statistical methodology was employed to compare OSCCs and controls. The quality assessment measurement of the samples within the pooled datasets was assessed by a. correlation of expression of housekeeping genes (optimally between 0.7-0.95. Control and OSCC datasets that were not fulfilling this were eliminated from further studies; b. consistency of replications. The consistency of replications was assessed by modified-standard deviation (SD) of the log-transformed expression in each sample from the tissue-specific median (where outliers with z > 3.5 are not used for estimating median); SD < 0.1 means good quality, and SD > 0.3 means bad quality; hence dataset that had SD≥0.3 were not used for further studies.^[18]^ The false discovery rate (FDR) based on Benjamini-Hochberg was ≤0.05 and fold change was fixed at 2. The scatter plot identified the over-expressed and under-expressed genes in OSCC vs controls. The list of the over-expressed and under-expressed genes of OSCC as compared to controls were collated and studied.

### 2.3 Ferroptosis Related Genes (FRG) Annotation

From the results of comparison, the list of over and under-expressed genes in OSCC was searched for FRG. The details of FRG were collated from two sources. One was the hand curated, peer-reviewed “Ferrdb” database http://www.zhounan.org/ferrdb/index.html while the other was from KEGG pathway.^[13, 19]^ The FRG were collated as drivers (including validated/deduced or suspected) or suppressors (including validated/deduced or suspected) or markers of ferroptosis The database provides links to publications in Pubmed database that mentions the gene-ferroptosis relationship. That was used as the standard reference material. Any unreferenced gene-ferroptosis connection discussed in this manuscript could be explored in the Ferrdb. From the KEGG Pathway (https://www.genome.jp/kegg-bin/show_pathway?map=hsa04216), the genes associated for the complete pathway (canonical and non-canonical) were collated.^[6,13,19]^

The significant genes from section 2.2 were compared with the list of collated FRGs. The genes in section 2.2 that were identified as FRGs were separated, listed and classified as suppressor or drivers or markers of ferroptosis.

### 2.4 Pathway deduction

The pathway by which the significant FRGs identified in the utility section of the Ferrdb. The significant relationship was flagged and fitted into the pathway established by the KEGG.

### 2.5 Validation of results

An attempt was made to validate the results using another OSCC and control cohort. The Human Cancer Genome Atlas(TCGA)-Head and Neck Squamous cell carcinoma (HNSCC) dataset (n=528) with control tissues(n=74) was accessed through the UCSC Xena browser (https://xenabrowser.net) and the genes identified studied for significance.^[20]^ The values of log^2^ (norm+1) was cast as Y axis. The Welsch t-test was used with P<0.05 as statistical significance.

## 3. RESULTS

### 3.1 Microarray datasets

From the publicly available database, in early May 2021, a total of 5 datasets (GSE 41613; GSE 37991; GSE56532; GSE64216; GSE30784) with 411 samples (316 OSCC and 95 controls) were collated as per the inclusion and exclusion criteria set were collected and processed in the Exatlas software as outlined. Of them, GSE 41613 had 97 OSCC; GSE 37991 had 40 OSCC and equal number of controls; GSE56532 had 10 OSCC and 2 controls; GSE64216 had 2 OSCC and 2 controls; GSE30784 had 167 OSCC and 45 controls. All these gene-sets were subjected to quality assessment measurement. Those sets whose house keeping genes did not fulfil the required minimum quality parameters of correlation or SD were removed from further analysis. After removal of non-qualifying datasets, a total of 259 OSCC (94 OSCC from GSE 41613; 165 from GSE 30784 and 44 controls from GSE 30784 were identified to form the final study population. Interestingly, all used GPL570 platform, ensuring further homogeneity in the pooled sample analysis.

### 3.2. Pairwise comparison

Pairwise comparisons were carried out and the scatterplot for the same are depicted in figure-1. All together there were 21139 genes with symbols and the significant genes (False detection rate, FDR≤0.05) were 12731. Of this only 1906 had fold change rate ≤2. Of this 1906 genes, 926 were overexpressed in OSCC and 980 under-expressed in OSCC as compared to controls.

**Figure-1:**
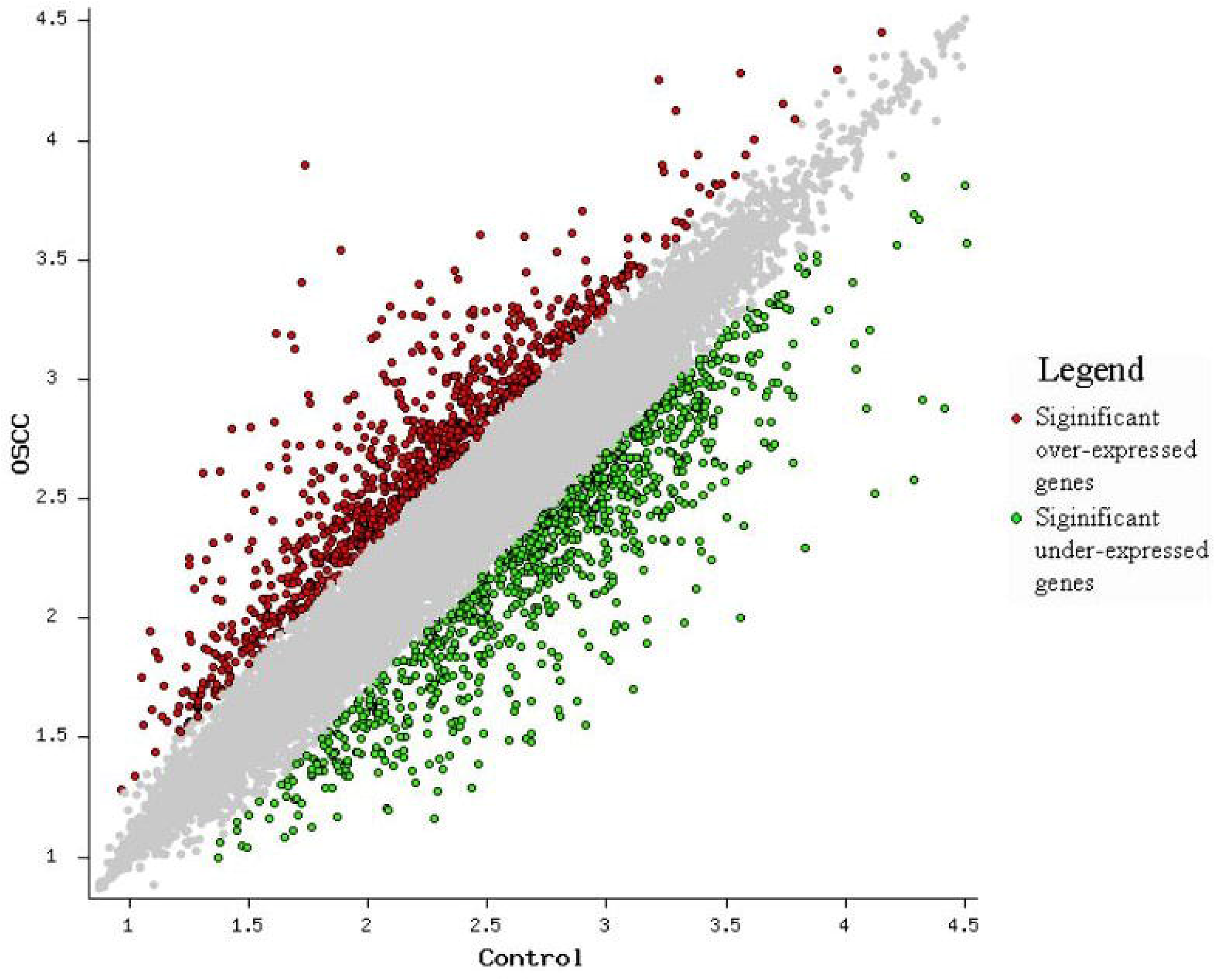
Scatter plot showing the differentially expressed genes between the Oral Squamous Cell Carcinoma (OSCC) and controls.

### 3.3 FRG Annotation

All the over and under-expressed genes were subjected to comparison with pooled list of FRGs. Eight genes from KEGG-ferroptosis pathway (CYBB, TFRC, TF, ACSL4, SLC7A11, CP, PRNP, SLC39A8 (ZIP8)) were identified. (Table-1)

**Table-1:** mRNA of Ferroptosis Related Genes (as identified from KEGG Pathway) differentially expressed between Oral Squamous Cell Carcinoma and Control tissues.

| Gene symbol | Experiment (@ P<0.001; FDR<0.001) |  |  |  | Validation |  |
| --- | --- | --- | --- | --- | --- | --- |
|  | OSCC | Control | Fold change | z- value | P-Value | T-statistics |
| SLC7A11 | 2.8631 | 2.4259 | 2.737 | 7.1170 | 0.007 | 3.59 |
| TFRC | 2.5130 | 2.1436 | 2.341 | 7.2630 | P<0.001 | 8.69 |
| CYBB | 2.4987 | 2.1649 | 2.157 | 7.6429 | 0.002 | 3.18 |
| ACSL4 | 2.6757 | 2.3682 | 2.030 | 10.7420 | 0.2970 | 1.05 |
| PRNP | 3.2048 | 2.9035 | 2.001 | 9.8939 | P<0.001 | 10.6 |
| SLC39A8 | 2.2980 | 1.9947 | 2.010 | 5.5739 | 0.78 | 0.26 |
| CP | 2.2957 | 2.7788 | 3.042 | 5.9333 | P<0.001 | -6.74 |
| TF | 2.1353 | 3.0122 | 7.532 | 8.0670 | P<0.001 | -14.05 |

There were 259 FRGs in the Ferroptosis database. Of this, in the present study, 44 (~17%) FRGs were identified. Of the 44, 21 were that promoted ferroptosis. Of this, 17 were validated drivers of ferroptosis (TF, ACSL4, ALOX12, BID, TNFAIP3, LPIN1, PRKAA2, MYB, PGD, CDO1, ALOX15B, ALOXE3, SOCS1, ALOX12B, PANX1, NOX4, CDKN2A) 3 deduced (CYBB, DUOX1, DUOX2) and 1 screened (TFRC). Of the 21 FRGs that drives ferroptosis, 9 (LPIN1, PRKAA2, MYB, PGD, CDO1, ALOX12, DUOX1 and DUOX2) were found significantly elevated in controls while the remaining 12 (CYBB, TFRC, ACSL4, BID, TNFAIP3, ALOX15B, ALOXE3, SOCS1, ALOX12B, PANX1, NOX4, CDKN2A) were elevated in OSCC. (Table-2)

**Table 2:**
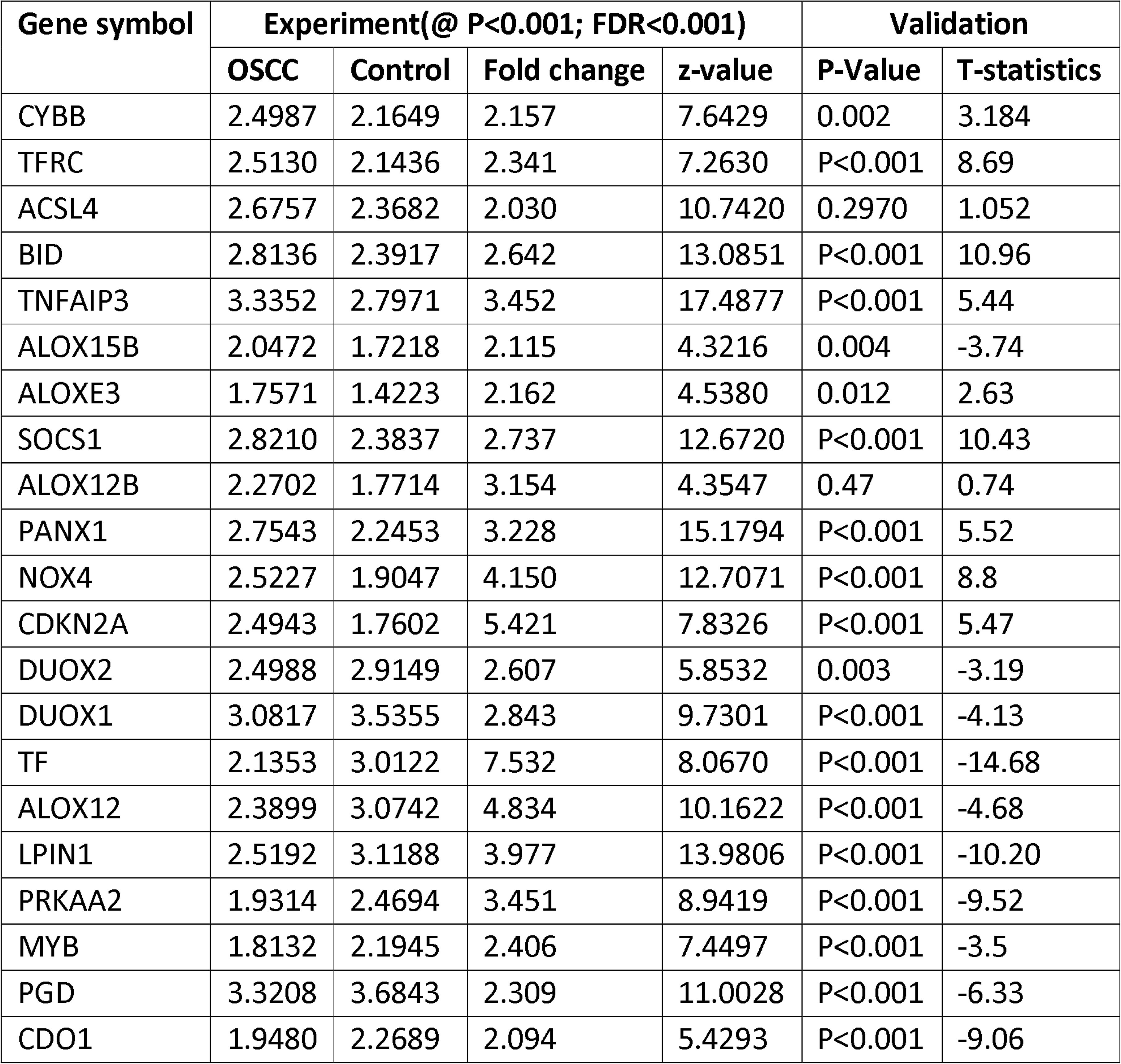
mRNA of drivers of Ferroptosis (as identified from ferroptosis database) differentially expressed between Oral Squamous Cell Carcinoma and Control tissues

There were 6 validated suppressors (SLC7A11, MUC1, AIFM2 (AKA FSP1), PML, CD44, CAV1). Of this, MUC1 and AIFM2(aka FSP) were increased in controls while SLC7A11, PML, CD44, CAV1 were known to be increased in OSCC. (Table-3)

**Table-3:** mRNA of suppressors of Ferroptosis (as identified from ferroptosis database) differentially expressed between Oral Squamous Cell Carcinoma and Control tissues

| Gene symbol | Experiment (@ P<0.001; FDR<0.001) |  |  |  | Validation |  |
| --- | --- | --- | --- | --- | --- | --- |
|  | OSCC | Control | Fold change | z-value | P-Value | T-statistics |
| SLC7A11 | 2.8631 | 2.4259 | 2.737 | 7.1170 | P<0.001 | 3.59 |
| PML | 2.7136 | 2.4064 | 2.029 | 9.8842 | P<0.001 | 7.69 |
| CD44 | 2.2958 | 1.9634 | 2.150 | 7.8632 | P<0.001 | 5.42 |
| CAV1 | 3.4098 | 3.0307 | 2.394 | 8.9699 | P<0.001 | 8.35 |
| MUC1 | 2.6726 | 3.2935 | 4.177 | 10.4357 | P<0.001 | -4.05 |
| AIFM2<br>(AKA FSP1) | 2.3834 | 2.8101 | 2.671 | 12.3004 | 0.84 | -0.20 |

There were 18 marker genes of ferroptosis. Of this, 6 were known ferroptosis inhibitors (SLC7A11, SLC2A12, GABPB1, SLC2A14, SLC2A3, RGS4) and 12 promoters (TFRC, TF, ALOX12, IL33, GPT2, HBA1, LURAP1L, PTGS2, NCF2, CXCL2, IL6, NNMT). OF the known ferroptosis inhibitor markers, SLC2A12 was alone increased in normal while the rest were increased in OSCC (SLC7A11, GABPB1, SLC2A14, SLC2A3, RGS4). Of the known ferroptosis promoter markers, TF, ALOX12, IL33, GPT2 and HBA1 were increased in controls while TFRC, LURAP1L, PTGS2, NCF2, CXCL2, IL6 and NNMT were raised in OSCC. (Table-4)

**Table-4:** mRNA of markers of Ferroptosis (as identified from ferroptosis database) differentially expressed between Oral Squamous Cell Carcinoma and Control tissues

| Gene symbol | Influence on Ferroptosis | Experiment (@ P<0.001; FDR<0.001) |  |  |  | Validation |  |
| --- | --- | --- | --- | --- | --- | --- | --- |
|  |  | OSCC | Control | Fold change | z-value | P-Value | T-statistics |
| SLC2A12 | Inhibitor | 2.3610 | 2.7433 | 2.412 | 7.8856 | 0.004 | -2.96 |
| SLC7A11 |  | 2.8631 | 2.4259 | 2.737 | 7.1170 | P<0.001 | 3.59 |
| GABPB1 |  | 2.5232 | 2.2014 | 2.098 | 10.2877 | P<0.001 | 10.15 |
| SLC2A14 |  | 2.3845 | 1.8573 | 3.367 | 11.1902 | 0.012 | 2.62 |
| SLC2A3 |  | 2.7827 | 2.2218 | 3.638 | 12.8455 | 0.01 | 2.7 |
| RGS4 |  | 2.5990 | 1.9843 | 4.118 | 8.4965 | P<0.001 | 10.91 |
| TF | Promoter | 2.1353 | 3.0122 | 7.532 | 8.0670 | P<0.001 | -14.05 |
| ALOX12 |  | 2.3899 | 3.0742 | 4.834 | 10.1622 | P<0.001 | -4.68 |
| IL33 |  | 2.3765 | 2.7386 | 2.302 | 5.8207 | P<0.001 | -14.18 |
| GPT2 |  | 2.5925 | 3.1979 | 4.031 | 13.1029 | P<0.001 | 17.14 |
| HBA1 |  | 3.3527 | 3.7419 | 2.450 | 6.5936 | 0.026 | -2.30 |
| TFRC |  | 2.5130 | 2.1436 | 2.341 | 7.2630 | P<0.001 | 8.69 |
| LURAP1L |  | 2.3752 | 1.9636 | 2.580 | 8.1975 | Missing in dataset |  |
| PTGS2 |  | 2.8169 | 2.3725 | 2.782 | 5.3109 | P<0.001 | 3.55 |
| NCF2 |  | 2.9061 | 2.4243 | 3.032 | 13.6832 | P<0.001 | 9.38 |
| CXCL2 |  | 3.0690 | 2.5236 | 3.511 | 10.6693 | P<0.001 | -3.57 |
| IL6 |  | 2.7053 | 2.0146 | 4.906 | 9.6193 | 0.011 | -2.66 |
| NNMT |  | 3.3090 | 2.6049 | 5.059 | 15.1900 | 0.99 | 0.005 |

### 3.4 Pathway deduction

Based on the utility aspect of the FRG database, FerrDB maintained online, in May 2021 the (http://www.zhounan.org/ferrdb/operations/buildregnet.html#) the pathway for the ferroptosis genes were collated. The same is given in figure-2.

**Figure-2:**
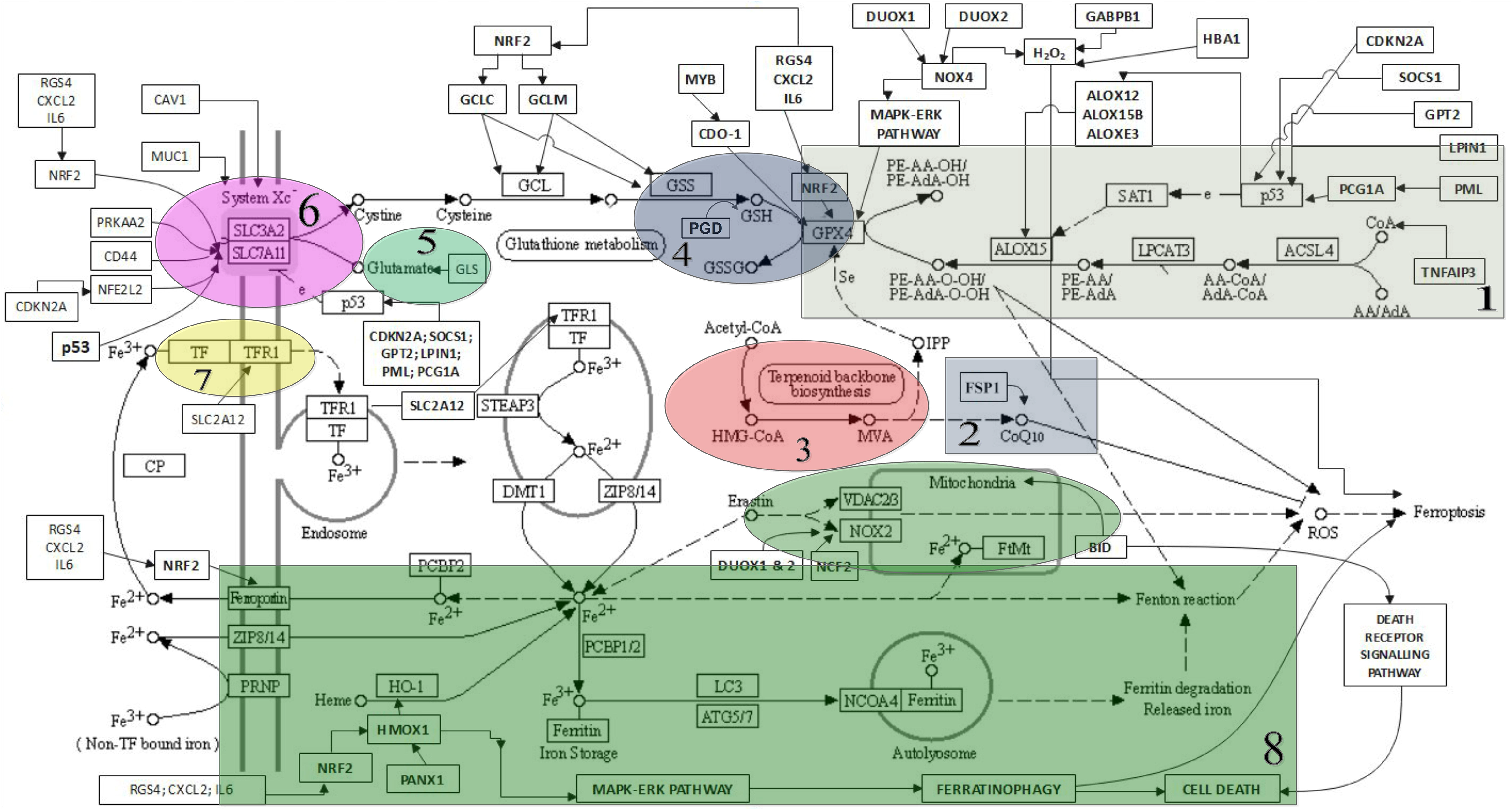
The Pathway of interaction of genes in Ferroptosis. 1. Lipid metabolism pathway; 2. FSP1-CoQ10-NADPH pathway; 3. Mevalonate pathway; 4. GPX4 pathway; 5. Glutaminolysis pathway; 6. Cystine deprivation-induced - System Xc pathways; 7. Iron metabolism pathway; 8. Ferritinophagy mechanism.

### 3.5 Validation of the results

An attempt was made to validate the results using the existing large scale OSCC dataset to identify the association of FRG expression identified in the present study. A comparison was made using the https://xenabrowser.net and TCGA HNSCC dataset of mRNA expression. The results of this are incorporated in Table-1 to 4 in validation column as well as in the figures 3, 4, 5 and 6.

**Figure-3:**
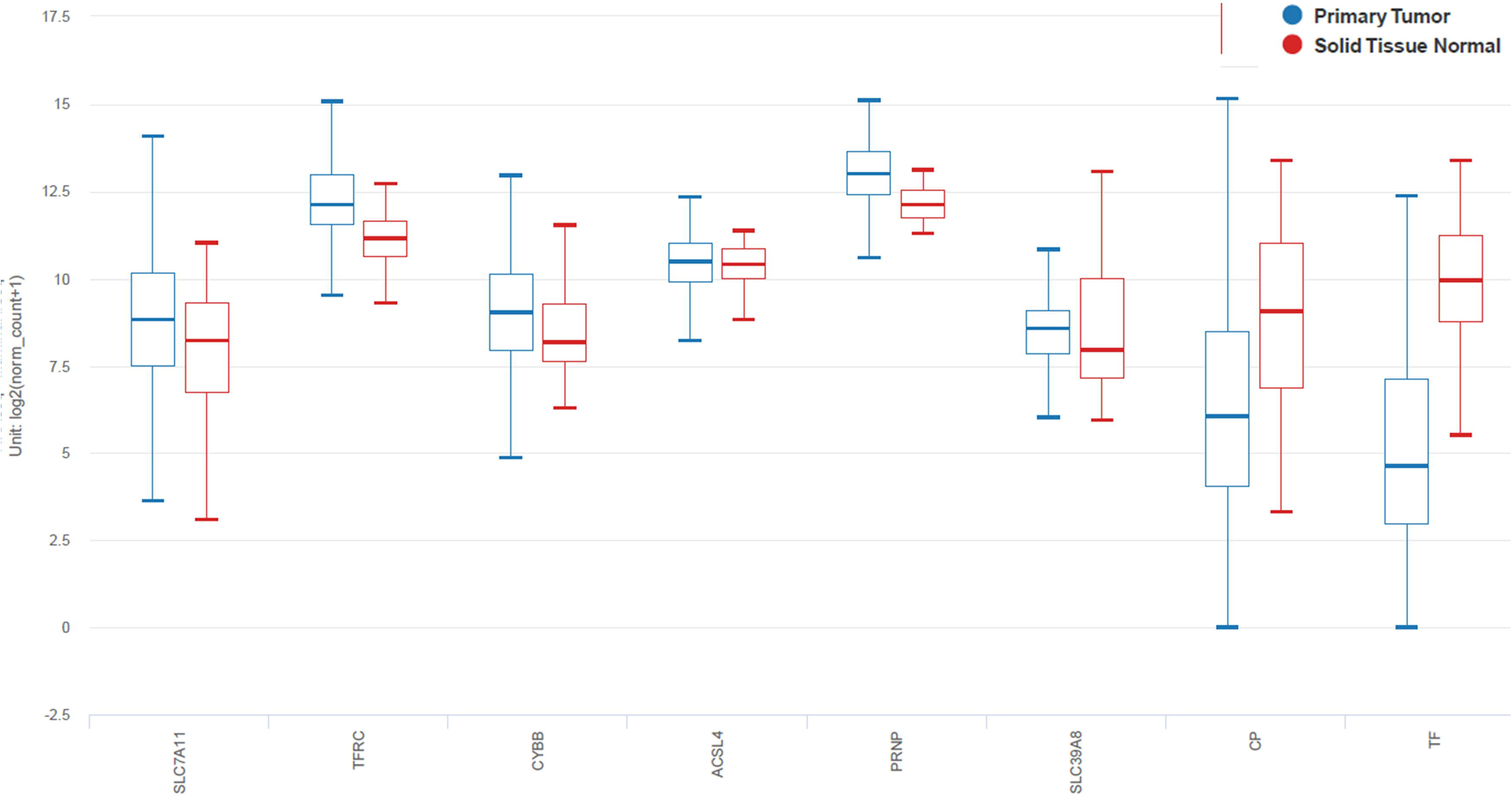
The validation study (with data from The Cancer Genome Atlas - Head and Neck Squamous Cell carcinoma) of KEGG Ferroptosis pathway.

**Figure-4:**
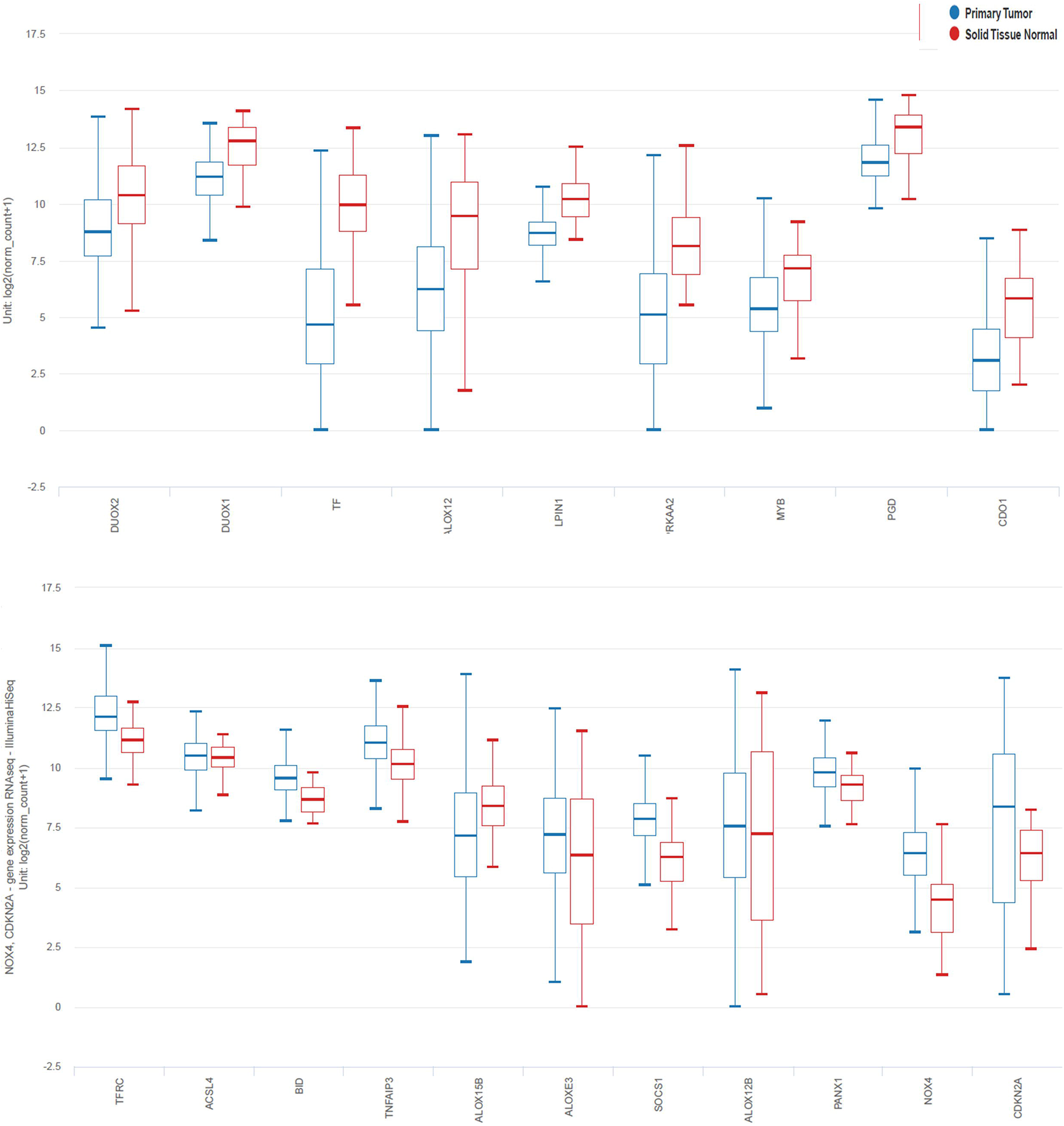
The validation study (with data from The Cancer Genome Atlas - Head and Neck Squamous Cell carcinoma) of drivers of ferroptosis. A. List of genes whose mRNA were increased in controls. B. List of genes whose mRNA were increased in Oral Squamous Cell Carcinoma (OSCC).

**Figure-5:**
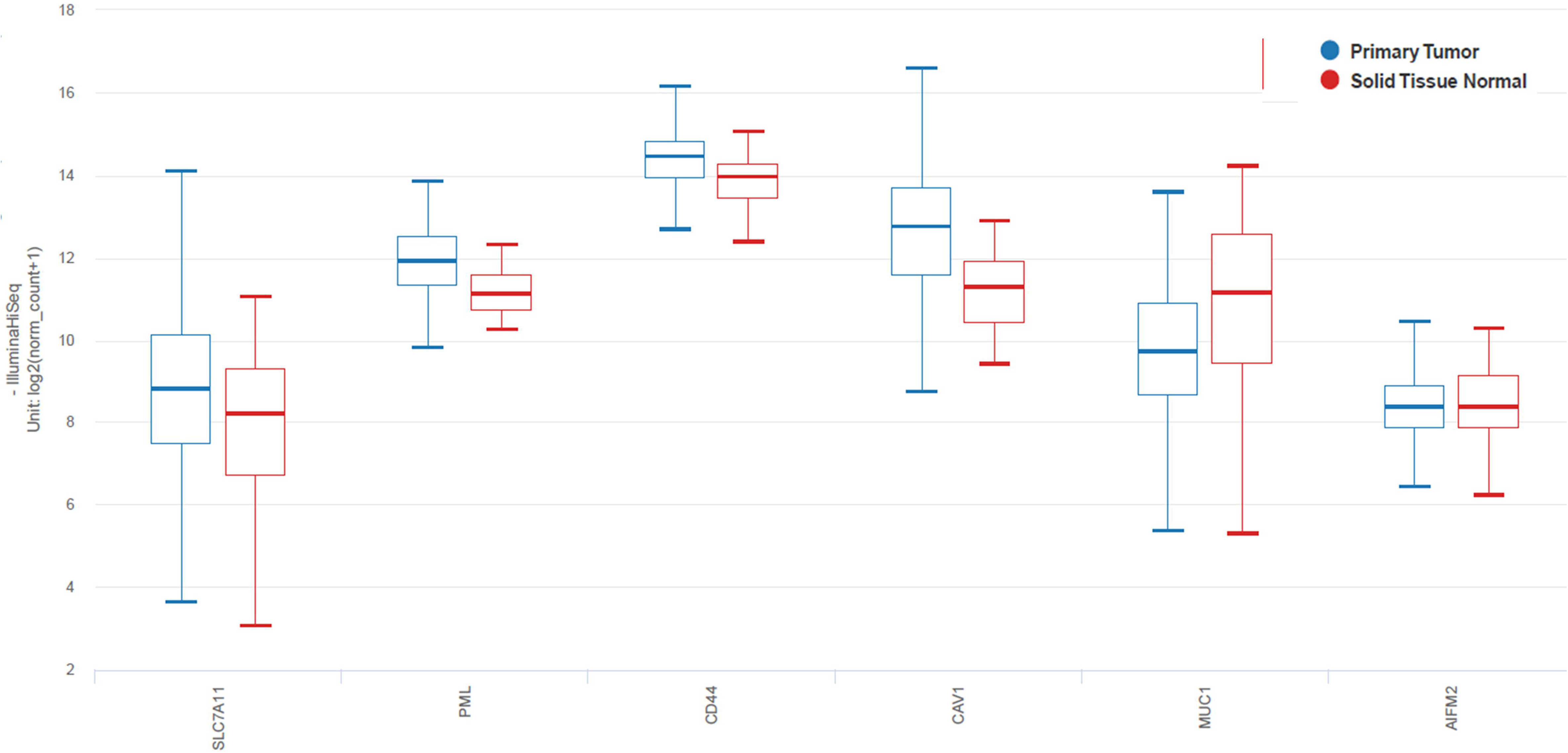
The validation study (with data from The Cancer Genome Atlas - Head and Neck Squamous Cell carcinoma) of suppressors of ferroptosis.

**Figure-6:**
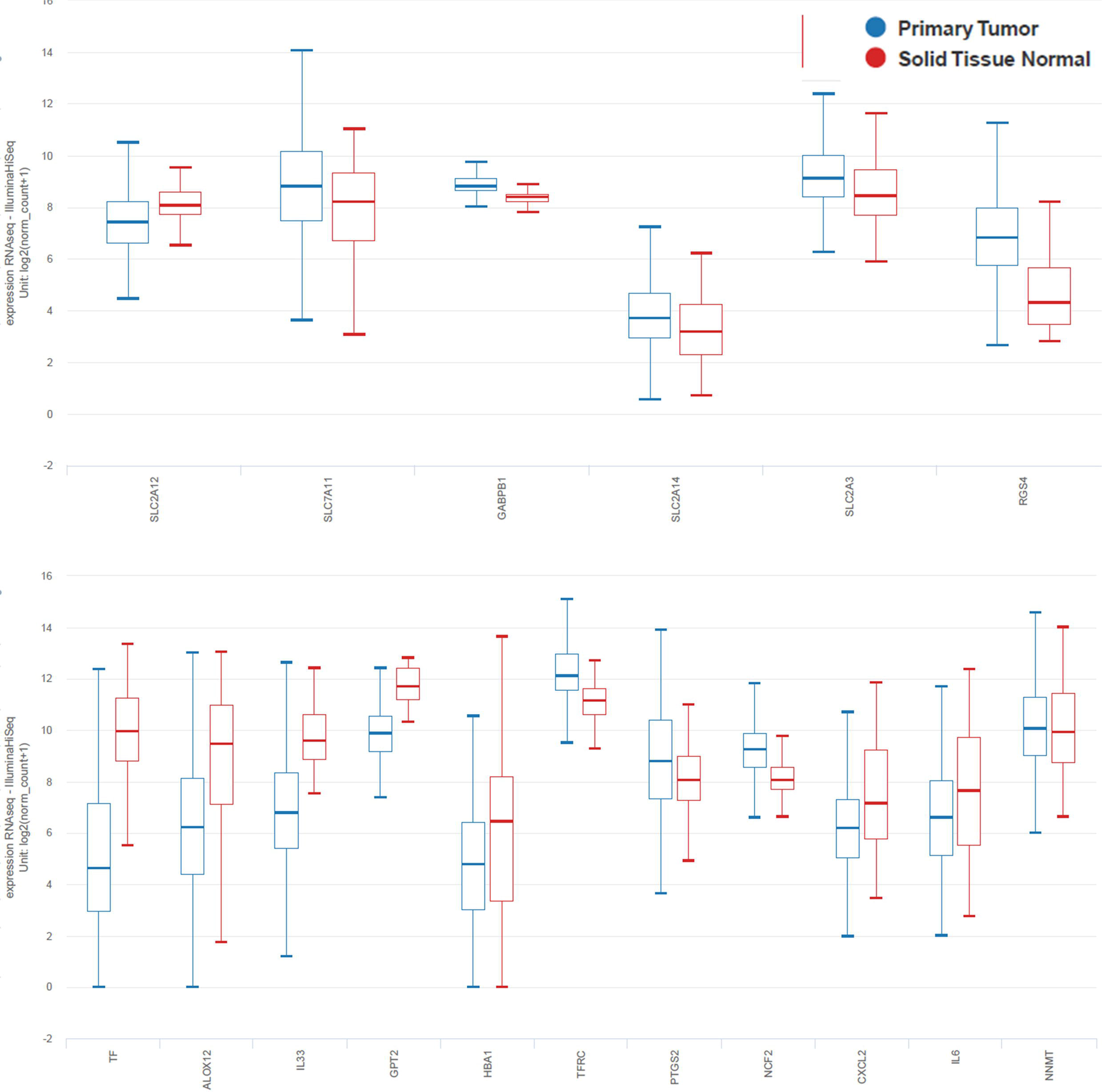
The validation study (with data from The Cancer Genome Atlas - Head and Neck Squamous Cell carcinoma) of markers of ferroptosis. A. Known ferroptosis inhibitors. B. Known ferroptosis promoters.

AIFM2 gene (Ferroptosis suppressor) and NNMT (Marker of ferroptosis – promoter) expression was in favour of controls, as in present study. However they were not statistically significant between OSCC and Controls in TCGA database. There was another discordancy limited to gene ALOX15B, CXCL2 and IL6. In present study, their higher expression was observed in OSCC while in TCGA dataset, they were high in controls. (Table-2, 4). The gene LPRIP1 was not obtained in the TCGA dataset. In all 5 genes were discordant with present results while one was unknown. Of the 6, 4 were associated with marker and one established driver and one suppressor gene had expression disparity.

## DISCUSSION

The neoplastic cells dynamically alters the expression of drivers/suppressors of ferroptosis to evade onset of ferroptosis and resultant death. This alters the lipid peroxidation, intracellular labile iron pool, enzymes and transport system associated with them. The evasion occurs with the suppressing of driver-promoter of ferroptosis or by over-expression of the suppressors of ferroptosis. Such a change is accompanied by alteration of ferroptosis markers.^[7–12, 21]^

There has been increasing interest in use of ferroptosis to selectively stimulate cell death in neoplastic cells.^[17]^ The association of FRG has not been widely described in OSCC literature. Hence an attempt was made to understand the association of ferroptosis in OSCC.

Of the 259 FRG, 44 genes (16.98%) were differentially expressed in OSCC compared to normal controls. SLC7A11 makes up the system X_c_^−^ that facilitates cysteine uptake into cell. SLC7A11 is a known suppressor of ferroptosis.^[6]^ Increased expression of this gene helps to evade ferroptosis. Increased expression of SLC7A11 is associated with poor prognosis of OSCC.^[22,23]^

Extracellularly, ceruloplasmin (CP) mediates the conversion of Fe^2+^ to Fe^3+^.^[24]^ This is crucial as most of the intracellular transport of iron is in Fe^2+^ form.^[6]^ Ceruloplasmin, is reduced in OSCC, due to attempt by neoplastic cells to evade accumulation of intracellular iron. Also, PRNP, a ferrireducatase that converts non-transferrin bound Fe^3+^ to Fe^2+^ is increased in OSCC.^[25]^ Solute Carrier 39A8 (SLC39A8 or ZIP8) is a transmembrane protein of plasma membrane and mitochondria, associated with transport of Fe^2+^ into the cell.^[6]^ This entity is increased in OSCC, which however is not validated.

The additive effect of CP, PRNP and SLC39A8 can reduce the intracellular transport of Fe^3+^ ions. The non-canonical, non-enzymatic iron transport pathway- TF receptor, TFRC or the TFR1 increases intra-cellular iron, facilitating ferroptosis.^[6]^ The expression of this gene is also increased in the report and validation. However, the levels of transferritin(TF), which works with TFRC, both at cell membrane and in endosomes are decreased both in the study and validation. It is possible that by modulating TF and TFRC together as a functional unit, the neoplastic cells try to evade ferroptosis by altering intracellular iron. Increase of intracellular Fe^2+^ would increase Fenton reaction. The ferroportin gene, SLC40A1 is the only transmembrane protein that transports Fe^2+^ from intracellular to extra-cellular spaces and also affects its reutilization.^[26]^ This process is influenced by RGS4, CXCL2 and IL6 through NRF2, a known redox sensitive transcription factor.^[26]^ This nuclear factor erythroid 2-like 2 (NFE2L2/NRF2) is a key transcription entity that plays a role in cell survival amidst oxidative stress by triggering the detoxification and antioxidant genes. Many NFE2L2 targeted genes are upregulated in ferroptosis and these genes are involved in iron metabolism (FTH1, SLC40A1, HMOX1 and GSH metabolism (SLC7A11).^[28]^ In the present study, ferroptosis inhibitor RGS4 was upregulated in OSCC as compared to controls, probably to evade ferroptosis. Ferroptosis promoters CXCL2 and IL6 were increased in OSCC. The RGS4, CXCL2 and IL6 also through the NRF2 activates the HMOX1, PANX1, HO1 can increase intracellular Fe^2+^ pool, facilitating ferratinophagy. The MAPK/ERF pathway is a signalling pathway that controls basic cellular processes and often is altered in the neoplastic process.^[27,28]^ In the present study PANX1 and IL6, driver-promoters of ferroptosis are elevated in OSCC. This pathway also act on GPX4.^[29]^ DUOX1, DUOX2 that are drivers of ferroptosis are reduced in the OSCC while NOX4, another ferroptosis driver is increased in OSCC. The ferroptosis inhibitor GABPB1 is increased in OSCC. This can increase hydrogen peroxide and act with NOX4. This influences the MAPK/ERF pathway and possibly acts on GPX4.

In the mitochondria, NOX2 is influenced by DUOX1, 2 and NCF2.^[6]^ DUOX1, DUOX2 are known drivers of ferroptosis and their expressions were identified to be reduced in OSCC as compared to controls and was validated. However, the NCF2, another potential marker of promoter of ferroptosis was elevated in OSCC than controls. BID is mediator of mitochondrial damage (via caspase-8).^[30]^ It is a known driver of ferroptosis and is elevated in OSCC than controls in the present study. This interplay of promoter-suppresser of ferroptosis along the mitochondria could influence the death receptor signalling pathway leading to cell death.

TNFAIP3 gene product is a known positive regulator of ACSL4 and a driver of ferroptosis.^[31]^ This gene is elevated in OSCC compared to control in the present study and also validated. The ACSL4-ALOX-15-ALOX12B-ALOX12, a part of the enzymatic non-canonical ferroptosis pathway had opposing expressions in the present study. The ferroptosis drivers ACSL4, ALOX15B, ALOXE3, ALOX12B were elevated while the ALOX12 was reduced. The inverse relationship of SLC7A11 and ALOX12 in driving ferroptosis is already known.^[32]^ Together, it is possible that OSCC cells attempts to evade ferroptosis by increasing suppressor SLC7A11 and suppressing the promoter ALOX12.

The role of p53 varies by the pathway it interacts with FRGs.^[6,14]^ It behaves as a pro-death entity when it interacts with SLC7A11, SAT1 and GLS2. It acts as a pro-survival molecule/gene when it acts via DPP4 and CDKN1A.^[6,14]^ In the present study, except SLC7A11, none of the known inter-actors were differentially expressed between OSCC and controls. However, CDKN2A was elevated in OSCC. CDKN2A is a known driver of ferroptosis that acts via the p19ARE pathway and through p16 with RB1 and p53 pathways.^[6,33–35]^ The increase of CDKN2A could thus influence the p53 pathway in neoplastic propagation.^[14]^ In addition, this study identified p53 interacters such as SOCS1, GPT2, LPIN1, GPT2 and PML. SOCS1 and GPT2 are known drivers of ferroptosis that are increased in OSCC in present study while GPT2 and LPIN1, both promoters are reduced in OSCC. PML is a known ferroptosis suppressor that is elevated in OSCC in this study.^[6,36]^ This interplay of p53 influenced drivers-suppressors of ferroptosis could variably influence the outcome.

Depending on the degree of expression of these genes and proteins, at the end, a neoplastic cell could either evade or succumb to ferroptosis.

In the non-enzymatic, non-canonical pathway, the FSP1 (aka AIFM2) in plasma membrane reduces coenzymeQ10 to ubiquinol that blocks lipid peroxidation. Reduction of FSP1, a known ferroptotic suppressor would facilitate neoplastic cell to evade PCD.^[6]^ In the present study, FSP1 is reduced and could not be validated in the TCGA database.

Cyclooxygenase- prostaglandin-E2 inflammation pathway is associated with OSCC and Prostaglandin-endoperoxide synthase(PTGS) is a key enzyme in prostaglandin biosynthesis.^[37]^ An inducible isoenzyme, PTGS2 encodes for cyclooxygenase-2(COX-2). It is reported that upregulated PTGS2 indicates ferroptosis onset and does not affect ferroptosis development.^[38]^ The elevated PTGS2 in OSCC as compared to controls and its subsequent validation in the present study indicate that ferroptosis happens in OSCC.

Genes encoding for CXCL2, NNMT and IL33 are elevated in OSCC in the present study. CXCL2 encodes for proteins related to immune regulation and neutrophils. Its overexpression has been observed in several tumors including gastric and renal carcinomas. Its elevation is associated with metastasis.^[39,40]^ Nicotinamide N-Methyltransferase or the NNMT is a metabolic regulator. It is altered and associated with a worse prognosis in various cancers. It is reported to promote epithelial-mesenchymal transition by modulating the transforming growth factor-β1. It is a pro-ferroptotic factor in a GSH-dependent manner. IL-33 elevation and its ferroptotic association has been observed in respiratory and urinary pathologies.^[41]^

In tumors, it has been reported that ferroptosis has a dual role of tumor promotion and suppression. The result is dependent on the gene, molecular patterns and pathways altered during the different phases of tumor initiation, growth and metastasis. Also, the damage-associated molecular patterns and the immune response triggered by FRGs damage within the tumour microenvironment contribute to the result.^[11]^ In the present study, it was observed that commonly the non-canonical pathway genes are more differentially expressed, both in pro-ferroptotic and anti-ferroptotic pathways. Recently, the role of FRGs in 20 cancers, including HNSCC has been described. The report indicates of mutations and alterations in HNSCC with high ferroptosis potential index highlighting the impact of the potential alteration of FRGs in HNSCC including OSCC.^[42]^

This study was designed with stringent, systematic, following all relevant recommended methods employed at every stage. Employing TCGA to validate the study further confirms the findings of the present study. However, TCGA has own limitations. For HNSCC, there is a relatively less number of cases with adjacent normal tissues in the TCGA database.^[43]^ The field-cancerization effect in the adjacent, apparently normal tissues at the gene and microRNA level can affect the findings.^[44]^ The normal tissue adjacent to tumor also has been demonstrated to be at an intermediate state between health and tumor states.^[45]^ Also, the TCGA database, in HNSCC includes oro-pharyngeal SCC lesions that might partially contribute to non-validation of site specific genes. Another fact that needs to be considered is that the bulk of the tumour would be from the epithelial component while in normal tissues, sizable entity would be from the connective tissue also.^[43]^ This may cause proportional variations. The TGCA validation should be interpreted accordingly. Also, the present study relies on mRNA-level quantifications for FRGs, whereas the ferroptosis process is dependent on proteins and the correlation may not be necessarily synchronous or proportional.^[42]^

Association of FRGs with OSCC is confirmed by the current study. Future research works in this direction needs to focus on confirmation of the type of ferroptotic pathways and their determinants. It also should focus on the epithelial and mesenchymal compartments individually. The role of the non-canonical ferroptotic pathway alterations, particularly the protein component and its impact on the various stages of OSCC need to be studied to arrive at OSCC specific and efficient therapeutics.

## CONCLUSION

The differentially expressed FRGs in a cohort of OSCC has been studied and validated. The differentially expressed FRGs in OSCC as compared to normal tissues can alter the various complex metabolic, inflammation and other cellular pathways. This could have a bearing impact on the efficacy of chemotherapy, radiotherapy and immunotherapy of OSCC.^[45]^ Also pro-ferroptotic drugs could target and increase PCD, improving the treatment outcomes. Large scale studies are needed at protein level to ascertain the extent of ferroptosis and the influence of socio-biological-demographical factors on such FRG and protein expression.

## Authors Declare

NO CONFLICT OF INTEREST

## Notes

### Competing Interest Statement

The authors have declared no competing interest.

